# The endosomal trafficking protein Myopic differentially regulates Drosophila C virus infection in cultured cells and adult flies

**DOI:** 10.64898/2026.07.29.741437

**Authors:** Jiaxin Liu, Kexin Li, Qinyi Liang, Yunfan Huang, Xuye Yuan, Tatsuhiko Kadowaki

## Abstract

Positive-sense RNA viruses extensively exploit host membrane trafficking pathways to establish intracellular replication organelles required for efficient viral replication. However, the contribution of endosomal trafficking proteins to infection by Drosophila C virus (DCV), a natural dicistrovirus of *Drosophila melanogaster*, remains poorly understood. Here, we investigated the role of Myopic (Mop), a conserved ESCRT-associated endosomal trafficking protein, during DCV infection. Knockdown of *mop* in cultured S2 cells significantly increased DCV RNA and viral protein accumulation without affecting viral binding or entry, indicating that Mop restricts DCV replication at a post-entry stage. In addition, depletion of *mop* altered the intracellular distribution of viral proteins, suggesting that Mop may influence the organization of DCV-associated intracellular membrane compartments. Unexpectedly, fat body-specific knockdown of *mop* in adult flies produced the opposite phenotype, resulting in reduced viral accumulation and prolonged survival following systemic DCV infection. This antiviral phenotype was not associated with enhanced activation of the STING or JAK–STAT pathways. Instead, transcriptomic and proteomic analyses revealed extensive remodeling of gene and protein expression, including altered abundance of proteins associated with autophagy and RNA interference. Together, our findings identify Mop as a previously unrecognized regulator of DCV infection and reveal distinct cell-autonomous and systemic functions of this conserved trafficking protein. These results highlight the importance of endosomal trafficking in antiviral defense and demonstrate that host membrane trafficking factors can exert fundamentally different effects on virus infection at the cellular and organismal levels.

## Introduction

Viruses depend extensively on host cellular machinery to complete their replication cycles. Positive-sense single-stranded RNA viruses, in particular, exploit host membrane trafficking pathways to establish intracellular replication organelles that support efficient viral RNA synthesis while shielding viral replication intermediates from host immune surveillance (Wolff, Melia et al. 2020). Members of the family Dicistroviridae, including Drosophila C virus (DCV), remodel intracellular membranes to generate replication compartments following entry into host cells (Cherry, Kunte et al. 2006). Consequently, host proteins involved in endosomal trafficking, membrane remodeling and vesicular transport have emerged as important determinants of viral infection. Defining how these host pathways regulate virus replication not only improves our understanding of virus-host interactions but also provides insight into fundamental aspects of membrane biology.

DCV is one of the best-characterized natural viral pathogens of *Drosophila melanogaster* and has become an important model for investigating antiviral immunity in insects (Mondotte and Saleh 2018). Following systemic infection, DCV replicates in multiple tissues and causes extensive pathology, leading to reduced locomotor activity, intestinal dysfunction and premature death (Chtarbanova, Lamiable et al. 2014, Segrist, Miller et al. 2024). Previous studies have established that the RNA interference (RNAi) pathway constitutes the major antiviral defense against DCV, while additional signaling pathways, including Toll, JAK–STAT and STING, also contribute to host resistance (Dostert, Jouanguy et al. 2005, Galiana-Arnoux, Dostert et al. 2006, van Rij, Saleh et al. 2006, Ferreira, Naylor et al. 2014, Goto, Okado et al. 2018). Despite these advances, the cellular mechanisms that regulate DCV replication after viral entry remain incompletely understood, particularly the contribution of intracellular membrane trafficking to the viral life cycle.

The *Drosophila* gene myopic (mop) encodes the orthologue of mammalian HD-PTP (PTPN23), a Bro1 domain-containing protein that functions as an accessory component of the endosomal sorting complexes required for transport (ESCRT) machinery. Mop regulates multiple aspects of endosomal trafficking, including receptor sorting, endosome maturation and lysosomal homeostasis, and is required for normal development and tissue homeostasis (Miura, Roignant et al. 2008, Chen, Li et al. 2012, Pradhan-Sundd and Verheyen 2015). In addition to its trafficking function, Mop has been implicated in innate immunity, where it is required for efficient Toll signaling and antimicrobial peptide induction (Huang, Chen et al. 2010). Because positive-strand RNA viruses rely heavily on host membrane remodeling and because innate immune signaling is intimately linked to endosomal trafficking, Mop represents an attractive candidate host factor that may influence DCV infection. However, whether Mop participates in antiviral defense or supports the DCV life cycle has not previously been investigated.

In the present study, we examined the role of Mop during DCV infection in cultured S2 cells and adult flies. We demonstrate that depletion of *mop* has strikingly different consequences in these two experimental systems. Knockdown of *mop* in S2 cells enhances DCV replication without affecting viral binding or entry, indicating that Mop functions as a cell-autonomous restriction factor during the post-entry stage of infection. Unexpectedly, fat body-specific knockdown of *mop* in adult flies suppresses systemic DCV infection and prolongs host survival. Transcriptomic and proteomic analyses further identify altered endolysosomal homeostasis together with changes in autophagy- and RNAi-associated proteins as candidate mechanisms underlying this phenotype. These findings reveal a previously unrecognized dual role of Mop in antiviral defense and highlight the distinct contributions of intracellular membrane trafficking and systemic physiological regulation during viral infection.

## Materials and methods

### Virus stock and titration

The DCV stock, kindly provided by Dr. Tao Peng (Guangzhou Medical University), was propagated in *Drosophila* S2 cells maintained at 25 °C in Schneider’s medium supplemented with 10% fetal bovine serum, 50 U/mL penicillin, and 50 μg/mL streptomycin (Beyotime).

Viral titers were determined by endpoint dilution assay in 96-well plates. S2 cells were seeded into flat-bottom 96-well plates at a density of 3 × 10^4^ cells per well in 75 μL of culture medium. On the following day, 180 μL of culture medium was dispensed into each well of a round-bottom 96-well plate. A 10-fold serial dilution of the virus stock was prepared by adding 20 μL of virus suspension to the first well and subsequently transferring 20 μL into the next well until the twelfth dilution was reached. Subsequently, 25 μL of each viral dilution was added to four replicate wells containing S2 cells. After incubation at 25 °C for 2 days, the cells were fixed with 100 μL of 8% paraformaldehyde in PBS for 20 min. The fixed cells were permeabilized with PT (PBS containing 0.1% Triton X-100) for 5 min and then blocked with PT containing 5% bovine serum albumin (BSA) for 30 min. Rabbit anti-DCV VP1 antibody (1:3000 dilution) prepared in the blocking solution was added to each well, and the plates were incubated overnight at 4 °C. After three washes with PT (10 min each), the cells were incubated with horseradish peroxidase (HRP)-conjugated anti-rabbit IgG (Proteintech; 1:3000 dilution) in PT containing 5% normal goat serum for 3 h at 25 °C. Following three additional washes with PT, infected cells were visualized using the AEC Chromogen Kit (red) (Boster). Wells containing more than 10 VP1-positive cells were scored as positive for infection. Viral titers were expressed as the median tissue culture infectious dose (TCID_50_) and calculated using the Reed–Muench method.

### Generation of anti-DCV VP1 antibody

The C-terminal region of DCV VP1 (amino acids 167–265) was amplified by PCR using the primer pair DCV752-F-BamHI and DCV850-R-NotI. The PCR product was digested with BamHI and NotI, purified, and ligated into the corresponding restriction sites of the pGEX-6P-3 expression vector (Cytiva). The resulting plasmid was transformed into *Escherichia coli* BL21 cells. Transformed *E. coli* BL21 cells were cultured in 1 L of LB medium containing 1% glucose and 0.1 mg/mL ampicillin at 37 °C until the optical density at 600 nm reached approximately 0.5. The culture was then cooled on ice, and recombinant protein expression was induced with 0.1 mM isopropyl β-D-1-thiogalactopyranoside at 15 °C for 16 h.

Bacterial cells were harvested by centrifugation and resuspended in 100 mL of ice-cold TNED buffer (50 mM <u>T</u>ris-HCl, pH 8.0, 150 mM <u>N</u>aCl, 2 mM <u>E</u>DTA, and 1 mM <u>D</u>TT) supplemented with 0.5 % Triton X-100 and a protease inhibitor cocktail (Beyotime). Cell lysates were prepared by sonication using a Q700 Sonicator (Qsonica) at an amplitude of 100 on ice for 45 min (30-s pulses with 3- min intervals). After centrifugation, the supernatant was incubated with 1 mL of BeyoGold™ GST-tag Purification Resin (Beyotime) at 4 °C for 2 h with gentle rotation. The resin was subsequently washed five times with 10 mL of TNED buffer. To release the recombinant VP1 protein, 2 mL of TNED buffer containing 100 units of PreScission Protease (Beyotime) was added to the resin, followed by incubation overnight at 4 °C. The supernatant containing the cleaved VP1 protein was collected, and the resin was further eluted once with 2 mL of TNED buffer. The eluate was combined with the initial supernatant and dialyzed twice against 2 L of PBS containing 1 mM EDTA and 0.5 mM DTT at 4 °C for 24 h. The purified VP1 protein was then sent to GeneScript (Nanjing, China) for the production of rabbit polyclonal anti-DCV VP1 antibody.

### Preparation of dsRNA

DNA templates for *mop* dsRNA synthesis were generated by RT-PCR using total RNA isolated from S2 cells and primer pairs containing the T7 RNA polymerase promoter sequence at their 5′ ends (T7-mop-F/T7-mop-R and T7-mop#2-F/T7- mop#2-R). For GFP dsRNA synthesis, the primer pair T7-GFP-F and T7-GFP-R was used with a GFP-containing plasmid as the PCR template. Double-stranded RNAs (dsRNAs) were synthesized from the PCR-generated DNA templates using T7 RNA polymerase (Takara). Following *in vitro* transcription, complementary RNA strands were annealed to generate dsRNA. The sizes of both the PCR products and the synthesized dsRNAs were verified by agarose gel electrophoresis.

### Knock-down of *mop* in S2 cells

To knock-down *mop* in S2 cells, 3 × 10^4^ cells were incubated with 4 μg of dsRNA in 30 μL of serum-free medium for 1 h, followed by adding 170 μL of the complete medium, and then inoculated to 96-well plate. After incubating at 25 °C for 4 days, DCV was added to the cells at 0.1 TCID/cell. The cells were lysed after 24 h using 200 μL of TRIZOL (Sigma-Aldrich) for total RNA extraction or 100 μL of 1 × SDS-PAGE sample buffer (2 % SDS, 10 % glycerol, 10 % β-mercaptoethanol, 0.25 % bromophenol blue, 50 mM Tris-HCl, pH 6.8) for Western blot.

Total RNA was dissolved with 20 μL H_2_O, then 4 μL was used for 20 μL reverse transcription reaction using ReverTra Ace (TOYOBO) and random primers. Then, 1 μL of the reverse transcript was used for qPCR to quantify DCV genome RNA using DCV-qPCR-F and DCV-qPCR-R primers and *mop* mRNA using mop-qPCR-F and mop-qPCR-R primers. *D. melanogaster* 18S rRNA served as the reference gene, amplified with primers Dm18S-F and Dm18S-R. Relative DCV genome RNA and *mop* mRNA abundance was calculated using the ΔCt method, with one GFP dsRNA treated sample set to 1. Statistical comparison of DCV genome RNA and *mop* mRNA was conducted by Steel test (two-tailed) and Welch’s t-test (two-tailed), respectively.

For Western blot, the samples were heated at 95 °C for 5 minutes, and 20 μL was applied to two 10 % SDS-PAGE gels. One gel was stained with Instant Blue (Abcam) to verify equal protein loading, while proteins from the other were transferred to a nitrocellulose membrane (Pall Life Sciences). The membrane was blocked with 5 % BSA in PBST (PBS with 0.1 % Tween-20), incubated with rabbit anti-DCV VP1 polyclonal antibody (1:1000 dilution) overnight at 4 °C. After washing five times with PBST (for 5 minutes each), the membrane was incubated with IRDye 680RD donkey anti-rabbit IgG (H+L) secondary antibody (1:10,000 dilution, LI-COR Biosciences) in PBST containing 5 % skim milk at room temperature for 3 hours. After another round of washing, the membrane was visualized using ChemiDoc MP (BioRad). The band intensity of DCV VP1 was measured using Image-J and compared by Welch’s t-test (two-tailed).

### DCV binding and entry assays

DCV binding and entry assays were performed using control and *mop*- knockdown S2 cells prepared as described above in 96-well plates. To assess viral binding, the cells were preincubated at 4 °C for 15 min. The culture medium was then replaced with 100 μL of medium containing 4 × 10^6^ TCID_50_ of DCV that had been pre-equilibrated at 4 °C, followed by incubation at 4 °C for 30 min. After incubation, the cells were washed twice with 200 μL of ice-cold PBS containing 1 mM MgCl_2_ and 1 mM CaCl_2_, lysed with 200 μL of TRIzol reagent, and subjected to total RNA extraction.

To assess viral entry, cells were incubated with DCV-containing medium under the same conditions except that the incubation was performed at 25 °C for 2 h. The cells were then washed twice with PBS containing 1 mM MgCl_2_ and 1 mM CaCl_2_ at room temperature. Surface-bound virus was removed by treatment with 0.5 % trypsin for 10 min at room temperature, followed by two washes with 200 μL of ice-cold culture medium. Cells were subsequently lysed with 200 μL of TRIzol reagent for total RNA extraction. DCV genomic RNA and *D. melanogaster* 18S rRNA were quantified by RT-qPCR as described above. Statistical comparisons of DCV genomic RNA levels were performed using a two-tailed Welch’s *t*-test.

### Immunofluorescence analysis of DCV VP1 in infected S2 cells

Control and *mop*-knockdown S2 cells were infected with DCV as described above and cultured on poly-L-lysine-coated eight-well chamber slides. Immunofluorescence staining was performed using the same procedure as described for the viral titration assay, except that fluorescein isothiocyanate (FITC)-conjugated anti-rabbit IgG (Beyotime; 1:500 dilution) was used as the secondary antibody and incubated with the samples for 3 h at room temperature. Following washing with PBS, nuclei were counterstained with DAPI (1 μg/mL) for 5 min and briefly rinsed with PBS. Images were acquired using a Zeiss LSM880 laser-scanning confocal microscope. Raw confocal images were processed using Fiji software, including cropping, channel merging, brightness and contrast adjustment, and maximum-intensity projection of Z-stack images. Final figures were assembled using Adobe Photoshop by overlaying bright-field and fluorescence images.

### Knockdown of mop in D. melanogaster

Adult female flies carrying *mop* knockdown constructs were generated by crossing Yolk-GAL4 (Bloomington Drosophila Stock Center #58814) or Hsp70- GAL4 (Bloomington #1799) driver lines with either UAS-*mop* shRNA (Bloomington #32916) or UAS-*mop* dsRNA (Bloomington #28522) lines. All fly stocks were confirmed to be free of *Wolbachia* and viral infections and carried the virus-sensitive allele of *pastrel*.

For systemic virus infection, adult female progeny and the corresponding parental control flies were injected with 50 nL of DCV (50 TCID_50_) into the thorax using a microinjector. Twenty-four hours after infection, 50 flies from each genotype were divided into five groups of 10 flies. Each group was homogenized in 400 μL of TRIzol reagent for total RNA extraction. Purified RNA was dissolved in 100 μL of RNase-free water, and 3 μL was used for reverse transcription as described above. The resulting cDNA was diluted with 100 μL of RNase-free water prior to quantitative PCR analysis. The abundances of DCV genomic RNA, *Srg3* mRNA (using primer pair Srg3-qPCR-F/Srg3-qPCR-R), and *TotM* mRNA (using primer pair TotM-qPCR-F/TotM-qPCR-R) were quantified by qPCR as described above. Relative transcript abundances were calculated using the ΔCt method, with one DCV-injected Yolk-GAL4 or Hsp70-GAL4 control sample normalized to a value of 1. Statistical comparisons were performed using the two-tailed Steel test. For Western blot analysis, three independent groups of 10 DCV-infected flies were homogenized in 300 μL of 1× SDS-PAGE sample buffer. After heating at 95 °C for 5 min, the homogenates were centrifuged briefly, and 10 μL of the supernatant was loaded onto two 10% SDS-polyacrylamide gels. One gel was stained with Instant Blue, whereas proteins separated on the second gel were transferred to a nitrocellulose membrane and subjected to Western blot analysis as described above. The intensity of the DCV VP1 band was quantified using ImageJ software and compared using the two-tailed Dunnett’s test.

For survival analysis, three independent groups of 30 DCV-infected flies from each genotype were maintained in vials containing standard fly food at 25 °C. As injection controls, flies of the corresponding genotypes were injected with 50 nL of 10 mM Tris-HCl. The number of surviving flies was recorded daily. Survival curves were generated using the Kaplan–Meier method and compared using the log-rank (Mantel–Cox) test.

### Transcriptome and proteome analyses of the fat body of *mop*-knockdown *D. melanogaster*

Five independent groups of 15 DCV-infected flies were prepared as described above. At 24 h post-infection, abdominal fat bodies were dissected by removing the gut and ovaries. Total RNA was extracted from each group using 400 μL of TRIzol reagent and subjected to RNA sequencing (RNA-seq). All libraries were sequenced on the DNBSEQ-T7 platform (Beijing Genomics Institute), generating at least 6 GB of clean sequencing data for each sample.

The quality of the RNA-seq data was evaluated using FastQC (Brown, Pirrung et al. 2017), and low-quality reads were filtered using SOAPnuke (version 2.2.1). After trimming low-quality reads and removing reads containing more than 5 % adapter sequences or ambiguous (N) bases, more than 92 % of the reads were retained for subsequent analyses. Clean reads were mapped to the *D. melanogaster* reference genome using STAR, and coordinate-sorted BAM files were generated. Principal component analysis (PCA) of the 15 RNA-seq samples was performed using the gmodels package (version 2.18.1) in R to evaluate sample clustering and reproducibility among biological replicates.

For differential gene expression analysis, htseq-count (Anders, Pyl et al. 2015) from the HTSeq package was used to quantify reads overlapping annotated gene features. Raw read counts were analyzed in RStudio (version 1.2.1335) using the DESeq2 package (version 1.24.0) from Bioconductor (Robinson, McCarthy et al. 2010). Read counts were normalized using the shrinkage estimation method implemented in DESeq2 (Love, Huber et al. 2014), and genes with fewer than one count per million mapped reads were excluded from further analysis. The Benjamini-Hochberg procedure was applied to estimate the false discovery rate (FDR), and Log_2_ fold changes (FC) were calculated for all detected genes. Differentially expressed genes (DEGs) among the three pairwise comparisons (Yolk-GAL4 vs. UAS-*mop* shRNA, Yolk-GAL4 vs. Yolk-GAL4>UAS-*mop* shRNA, and UAS-*mop* shRNA vs. Yolk-GAL4>UAS-*mop* shRNA) were identified using a threshold of Q-value < 0.05.

Gene Ontology (GO) enrichment analysis of the 152 DEGs was performed using the clusterProfiler package (version 3.18.0) (Yu, Wang et al. 2012). The most specific enriched GO terms were selected using an FDR cutoff of < 0.05. All statistical analyses were performed in RStudio (version 1.2.1335).

The RNA-seq data have been deposited in the NCBI Sequence Read Archive under accession number PRJNA1498194.

For quantitative proteomic analysis, abdominal fat bodies prepared as described above were suspended in 300 μL of 8 M urea, sonicated, and centrifuged to remove insoluble material. A total of 150 μL of the supernatant, corresponding to approximately 150–200 μg of protein, was submitted to the Beijing Genomics Institute for label-free quantitative proteomic analysis using the data-independent acquisition (DIA) workflow. Peptide separation was performed using a Thermo UltiMate 3000 UHPLC system. Eluted peptides were ionized using a captive spray ionization nanoliter source and analyzed on a timsTOF Pro tandem mass spectrometer (Bruker Corporation) operating in DIA mode. Peptide and protein identification and quantification were achieved by deconvolution of the DIA data using a spectral library generated from data-dependent acquisition analysis.

The MSstats software package was used for intra-system error correction, normalization of protein abundance values, and downstream differential protein analysis. PCA was performed using the normalized protein abundance matrix generated after MSstats normalization to evaluate the overall similarity among biological replicates and experimental groups. The first two principal components were used to visualize sample clustering according to genotype. A total of 48,035–50,217 peptides and 5,863–5,968 proteins were quantified in individual samples. Protein abundance fold changes were calculated for predefined group comparisons based on the quantitative values obtained for each protein. Statistical significance was assessed using Welch’s *t*-test, and proteins with a fold change greater than 1.5 and a *P* value < 0.05 were considered differentially expressed. In addition, group-specific proteins were identified as proteins detected exclusively in one experimental group but completely absent in the comparison group. The mass spectrometry proteomics data have been deposited to the ProteomeXchange Consortium via the PRIDE (Perez-Riverol, Bandla et al. 2025) partner repository with the dataset identifier PXD081508. All primers used in this study are listed in Supplementary Dataset 1.

## Results

### Knockdown of *mop* enhances DCV infection in S2 cells

To investigate the role of *myopic* (*mop*) in DCV infection, we depleted *mop* in S2 cells by RNA interference and quantified viral replication by RT-qPCR.

Knockdown of *mop* significantly increased the accumulation of DCV genomic RNA compared with control cells treated with GFP dsRNA (Fig. 1A). A similar increase in viral RNA was observed using a second, non-overlapping dsRNA targeting a different region of the *mop* transcript (mop dsRNA #2), indicating that the phenotype was not caused by an off-target effect (Fig. 1A). Efficient knockdown of *mop* was confirmed by RT-qPCR (Fig. 1B). To determine whether increased viral RNA was accompanied by enhanced viral protein production, VP1 expression was examined by immunoblotting. Consistent with the RT-qPCR results, VP1 accumulated to a significantly higher level in *mop*-knockdown (mop KD) cells than in control cells (Fig. 1C, D). Together, these results demonstrate that depletion of *mop* enhances DCV infection in S2 cells, indicating that Mop functions as a cell-autonomous restriction factor against DCV. This finding is consistent with the reported requirement of Mop for Toll-dependent induction of antimicrobial peptide genes (Huang, Chen et al. 2010).

**Figure 1.**
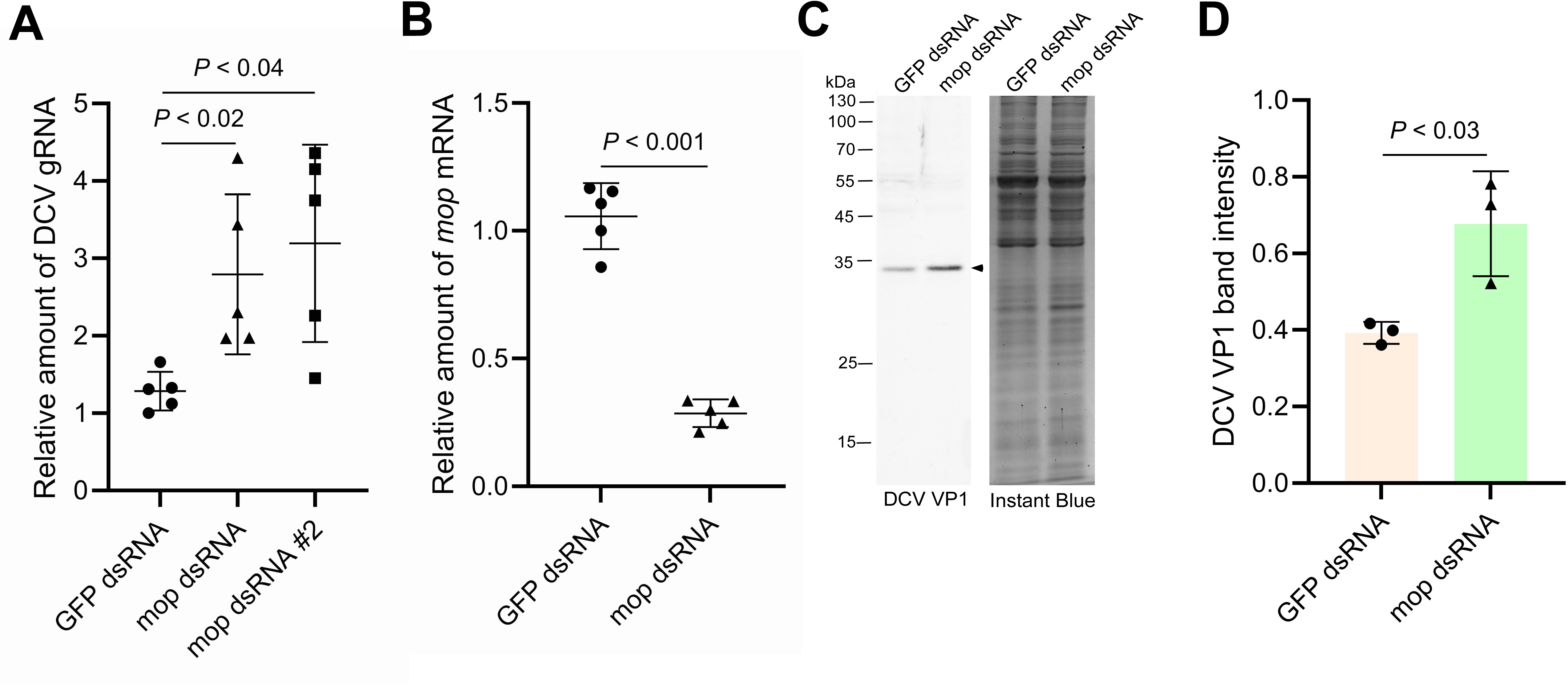
Knockdown of *mop* promotes DCV replication in S2 cells. (A) Relative abundance of DCV genomic RNA (DCV gRNA) in control (GFP dsRNA-treated) and *mop*-knockdown (mop dsRNA and mop dsRNA #2-treated) S2 cells at 24 h post-infection. Knockdown of *mop* significantly increased the accumulation of DCV gRNA (n = 5). (B) Relative abundance of *mop* mRNA in control (GFP dsRNA-treated) and *mop*-knockdown (mop dsRNA-treated) S2 cells. *mop* mRNA levels were significantly reduced following dsRNA treatment (n = 5). (C) Western blot analysis of DCV VP1 protein in lysates of DCV-infected control (GFP dsRNA-treated) and *mop*-knockdown (mop dsRNA-treated) S2 cells. Total protein was visualized by Instant Blue staining as a loading control. The DCV VP1 band is indicated by the arrowhead. Molecular weight markers (kDa) are shown on the left. (D) Quantification of DCV VP1 protein levels shown in panel (C). Knockdown of *mop* resulted in significantly greater accumulation of DCV VP1 compared with the control (n = 3).

### Knockdown of *mop* does not affect DCV binding or entry

To determine which stage of the viral life cycle is regulated by Mop, we first examined whether *mop* knockdown affected viral attachment or entry into S2 cells. Quantification of cell-associated virus revealed that both DCV binding and entry were comparable between control and mop KD S2 cells (Fig. 2A), indicating that Mop is not required for these early steps of infection.

**Figure 2.**
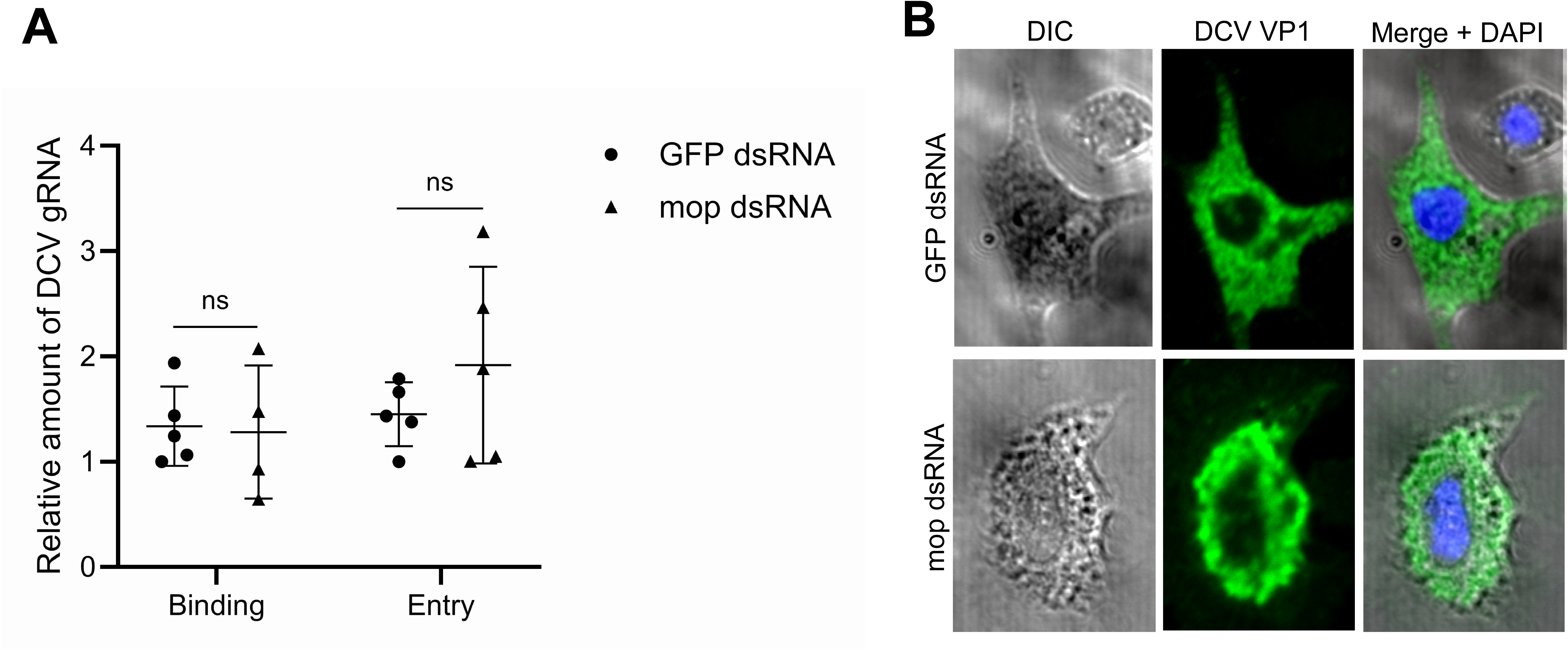
Mop affects DCV infection at a post-entry step. (A) Relative abundance of DCV bound to (30 min at 4 °C) or internalized into (2 h at 25 °C) control (GFP dsRNA-treated) and *mop*-knockdown (mop dsRNA-treated) S2 cells. No significant differences were detected in viral binding or entry between the two groups (n = 5, except *mop*-knockdown cells in the binding assay, n = 4). (B) Immunofluorescence analysis of DCV VP1 in DCV-infected control (GFP dsRNA-treated) and *mop*-knockdown (mop dsRNA-treated) S2 cells. Differential interference contrast (DIC) images and merged fluorescence images showing VP1 (green) and DAPI-stained nuclei (blue) are presented. In control cells, VP1 exhibited a dispersed cytoplasmic distribution, whereas in *mop*-knockdown cells, VP1 displayed a more punctate localization. Scale bar = 5 μm.

DCV replication is thought to occur on virus-induced membrane-associated replication organelles that depend on host membrane remodeling (Cherry, Kunte et al. 2006). Because Mop is an endosomal protein involved in endosomal sorting and vesicular trafficking (Chen, Li et al. 2012, Pradhan-Sundd and Verheyen 2015), we next examined the intracellular distribution of DCV in infected cells. In control S2 cells, VP1 staining was distributed broadly throughout the cytoplasm. In contrast, VP1 staining in mop KD cells appeared more punctate and was preferentially concentrated in the perinuclear region (Fig. 2B). These observations suggest that depletion of Mop alters the intracellular organization of DCV-containing compartments. This altered distribution is consistent with previous reports showing enlargement of endosomal compartments following loss of Mop function (Miura, Roignant et al. 2008).

### Knockdown of *mop* in adult fat body suppresses DCV infection

To investigate the role of Mop during DCV infection in vivo, *mop* was knocked down in adult flies using the GAL4/UAS system. Ubiquitous knockdown driven by Tubulin-GAL4 or Actin-GAL4 resulted in lethality, preventing further analysis. In contrast, flies carrying Yolk-GAL4, which drives expression predominantly in the adult female fat body, were viable following *mop* knockdown (hereafter referred to as mop KD flies).

Unexpectedly, systemic DCV infection was significantly reduced in mop KD flies compared with both parental control lines (UAS-*mop* shRNA and Yolk- GAL4), as determined by viral RNA levels (Fig. 3A). Similar results were obtained using an independent UAS-*mop* long dsRNA construct targeting a different region of the *mop* transcript, excluding the possibility of an off-target effect (Fig. 3A). Furthermore, knockdown of *mop* using Hsp70-GAL4 also resulted in reduced DCV infection (Fig. 3B), indicating that the antiviral phenotype is not specific to the Yolk-GAL4 driver. Consistent with the reduction in viral RNA, accumulation of VP1 protein was also markedly decreased in mop KD flies compared with the parental controls (Fig. 3C, D). Moreover, mop KD flies survived significantly longer following DCV infection than either parental line (Fig. 3E; log-rank test, χ² = 46.51, df = 2, *P* < 0.0001). Thus, unlike the phenotype observed in cultured S2 cells, depletion of *mop* in fat bodies of adult flies suppresses DCV infection and improves host survival.

**Figure 3.**
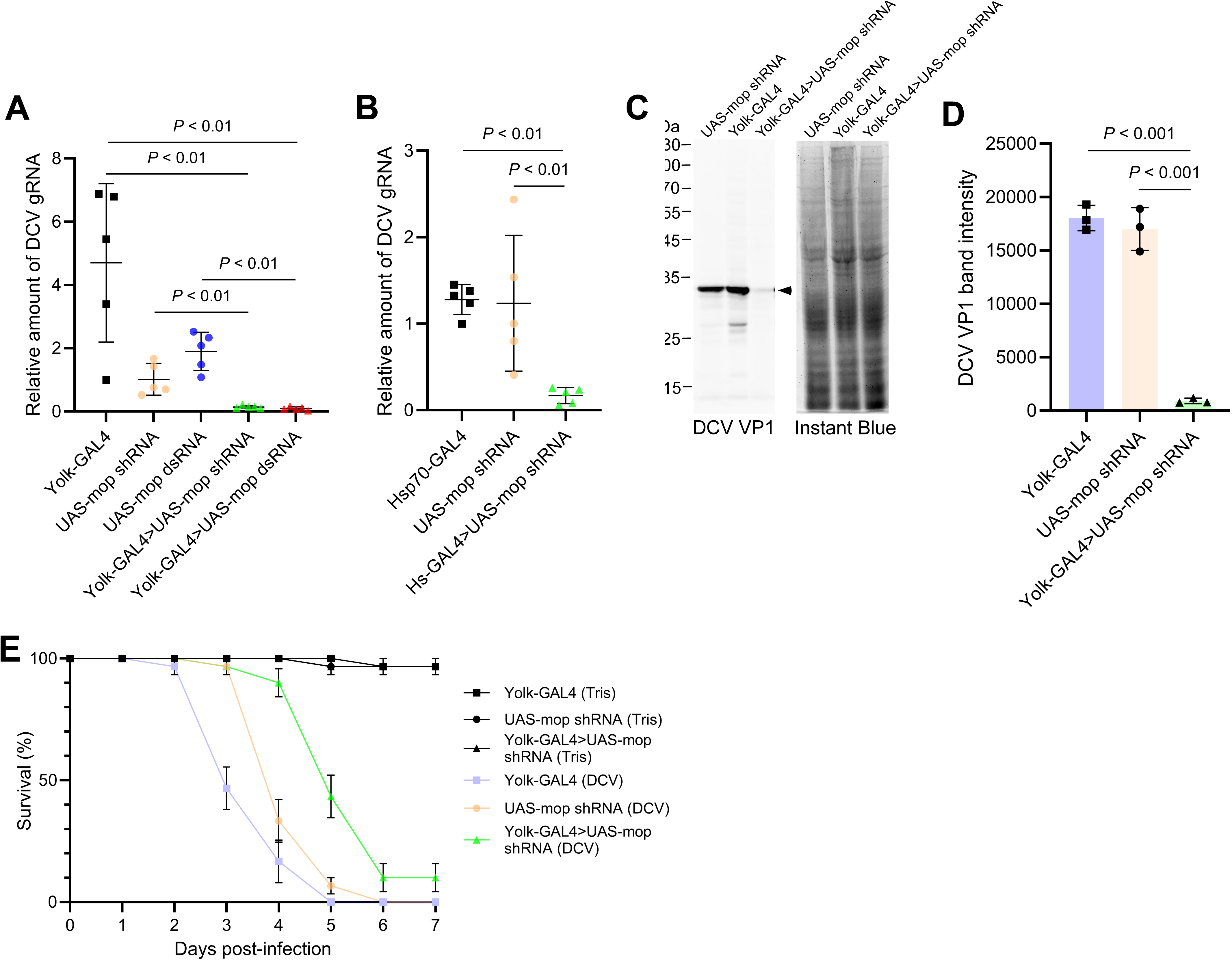
Knockdown of *mop* in the adult fat body suppresses DCV infection and improves survival. (A) Relative abundance of DCV genomic RNA (DCV gRNA) in parental control (Yolk-GAL4, UAS-mop shRNA, and UAS-mop dsRNA) and *mop*-knockdown (Yolk-GAL4>UAS-mop shRNA and Yolk-GAL4>UAS-mop dsRNA) flies at 24 h post-infection. Knockdown of *mop* in the adult fat body significantly reduced the accumulation of DCV gRNA (n = 5). (B) Relative abundance of DCV gRNA in parental control (Hsp70-GAL4 and UAS-mop shRNA) and *mop*-knockdown (Hsp70-GAL4>UAS-mop shRNA) flies at 24 h post-infection. Knockdown of *mop* using the Hsp70-GAL4 driver also significantly reduced DCV gRNA accumulation (n = 5). (C) Western blot analysis of DCV VP1 protein in lysates of DCV-infected parental control (Yolk-GAL4 and UAS-mop shRNA) and *mop*-knockdown (Yolk-GAL4>UAS-mop shRNA) flies. Total protein was visualized by Instant Blue staining as a loading control. The DCV VP1 band is indicated by the arrowhead. Molecular weight markers (kDa) are shown on the left. (D) Quantification of DCV VP1 protein levels shown in panel (C). *mop* knockdown significantly reduced the accumulation of DCV VP1 compared with the parental controls (n = 3). (E) Survival of parental control (Yolk-GAL4 and UAS-mop shRNA) and *mop*-knockdown (Yolk-GAL4>UAS-mop shRNA) flies following injection with either 10 mM Tris-HCl or DCV. Survival was monitored daily for 7 days after injection. *Mop* knockdown significantly improved the survival of DCV-infected flies (n = 3).

### Knockdown of *mop* in the adult fat body does not enhance STING- or JAK– STAT-dependent immune responses

The reduced viral burden observed in mop KD flies raised the possibility that depletion of *mop* enhances antiviral immune signaling. To examine this possibility, we measured the expression of *Srg3* and *TotM* mRNAs, representative downstream targets of the STING and JAK–STAT pathways, respectively (Dostert, Jouanguy et al. 2005, Goto, Okado et al. 2018). The abundance of *Srg3* mRNA was comparable between DCV-infected mop KD flies and UAS-*mop* shRNA control flies, although it was significantly lower in Yolk-GAL4 flies (Fig. 4A). In contrast, *TotM* mRNA was significantly reduced in mop KD flies compared with both parental controls (Fig. 4B). These results do not support enhanced activation of the STING pathway in mop KD flies. The reduced expression of *TotM* may instead reflect the lower level of DCV infection observed in these flies. Collectively, these data indicate that the decreased susceptibility of mop KD flies to DCV infection cannot be readily explained by increased activation of either the STING or JAK–STAT pathways.

**Figure 4.**
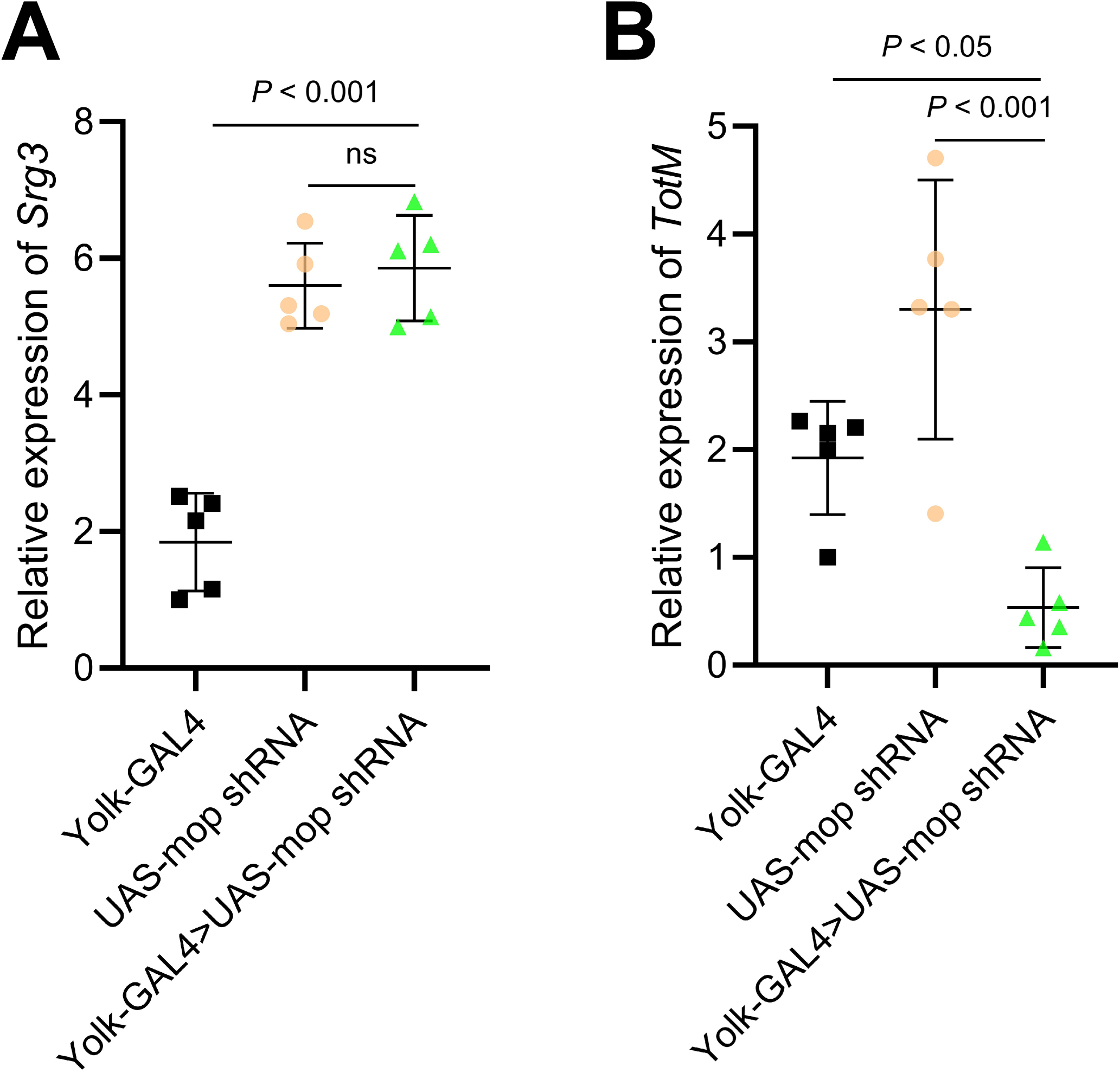
Expression of STING- and JAK–STAT-responsive genes in *mop*-knockdown flies. (A) Relative abundance of *Srg3* mRNA in parental control (Yolk-GAL4 and UAS-mop shRNA) and *mop*-knockdown (Yolk-GAL4>*UAS*-mop shRNA) flies at 24 h post-infection. *Srg3* mRNA levels were comparable between UAS-mop shRNA and *mop*-knockdown flies but were significantly lower in Yolk-GAL4 flies (n = 5). (B) Relative abundance of *TotM* mRNA in parental control (Yolk-GAL4 and UAS-mop shRNA) and *mop*-knockdown (Yolk-GAL4>UAS-mop shRNA) flies at 24 h post-infection. *TotM* mRNA levels were significantly lower in *mop*-knockdown flies than in either parental control (n = 5).

### Transcriptomic and proteomic characterization of mop KD flies

To obtain insight into the molecular basis underlying reduced DCV infection following fat body-specific *mop* knockdown, we performed transcriptomic and proteomic analyses of fat bodies isolated from DCV-infected mop KD flies and the corresponding parental controls. Principal component analysis (PCA) of the RNA-seq datasets showed clear clustering according to genotype (Fig. 5A), indicating that *mop* knockdown induces reproducible changes in gene expression. Differential expression analysis identified 152 transcripts whose abundance was specifically altered in mop KD flies relative to the parental controls, including 97 upregulated and 55 downregulated genes (Fig. 5B; Supplementary Dataset 2). Gene Ontology (GO) enrichment analysis of the upregulated genes revealed significant enrichment of the term cell-cell junction (Supplementary Dataset 3). In contrast, the downregulated genes were significantly enriched for the GO terms cellular response to UV, represented by *TotA*, *TotB*, and *TotC*, and ribosome biogenesis (Supplementary Dataset 3). The reduced abundance of *TotA*, *TotB*, and *TotC* transcripts is consistent with the decreased expression of *TotM* described above.

**Figure 5.**
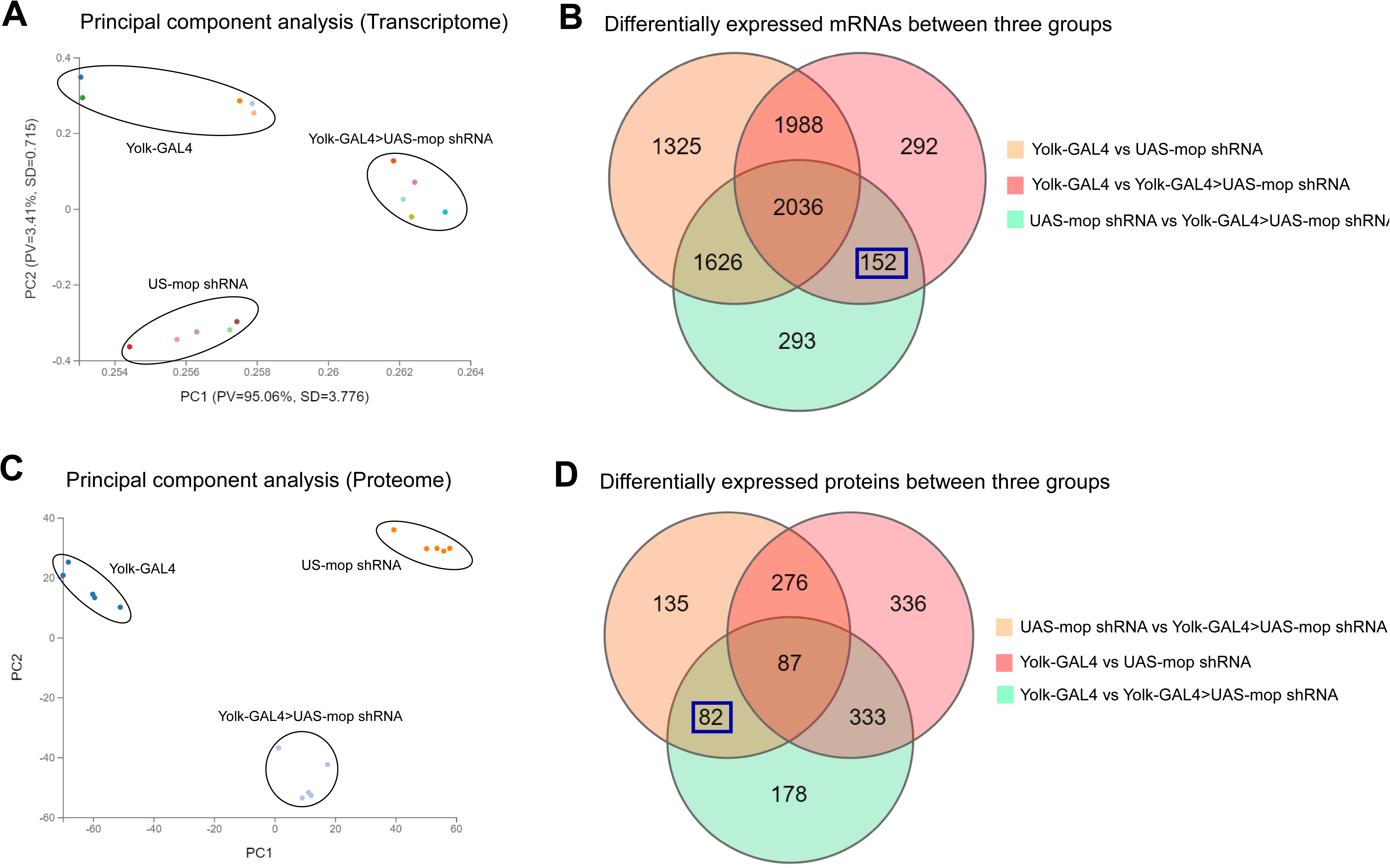
Transcriptomic and proteomic characterization of *mop*-knockdown flies. (A) Principal component analysis (PCA) of transcriptomes from parental control (Yolk-GAL4 and UAS-mop shRNA) and *mop*-knockdown (Yolk-GAL4>UAS*-mop* shRNA) flies at 24 h post-infection. Transcriptomic profiles clustered according to genotype. (B) Venn diagram showing differentially expressed mRNAs among parental control (Yolk-GAL4 and UAS-*mop* shRNA) and *mop*-knockdown (Yolk-GAL4>UAS-*mop* shRNA) flies. A total of 152 mRNAs were specifically up- or down-regulated in *mop*-knockdown flies relative to the parental controls (blue). (C) Principal component analysis (PCA) of proteomes from parental control (Yolk-GAL4 and UAS*-mop* shRNA) and *mop*-knockdown (Yolk-GAL4>UAS-mop shRNA) flies at 24 h post-infection. Proteomic profiles also clustered according to genotype. (D) Venn diagram showing differentially expressed proteins among parental control (Yolk-GAL4 and UAS-mop shRNA) and *mop*-knockdown (Yolk-GAL4>UAS-mop shRNA) flies. A total of 82 proteins were specifically up- or down-regulated in *mop*-knockdown flies relative to the parental controls (blue).

Proteomic analysis likewise demonstrated clear separation of samples according to genotype by PCA (Fig. 5C). A total of 82 proteins were differentially abundant in mop KD flies, including 44 upregulated and 37 downregulated proteins (Fig. 5D; Supplementary Dataset 4). Although no virus-related GO term was significantly enriched, two proteins with established antiviral functions, Ref(2)P and AGO2, accumulated in mop KD flies. Given the established role of Mop in endolysosomal trafficking, the increased abundance of Ref(2)P and AGO2 is consistent with altered endolysosomal homeostasis following *mop* depletion. Because Ref(2)P and AGO2 are key components of the autophagy and RNA interference pathways, respectively, their increased accumulation identifies these pathways as potential contributors to the reduced susceptibility of mop KD flies to DCV infection.

## Discussion

Our experiments in cultured S2 cells indicate that Mop acts at a post-entry step of the DCV life cycle. Viral attachment and entry were unaffected by *mop* knockdown, whereas viral RNA and VP1 protein accumulated to significantly higher levels. Moreover, DCV staining became more punctate and concentrated around the perinuclear region in mop KD cells. Positive-strand RNA viruses, including picorna-like viruses, replicate on membrane-bound replication organelles generated through extensive remodeling of host intracellular membranes (Cherry, Kunte et al. 2006, Wolff, Melia et al. 2020). Because Mop is an endosomal trafficking protein, these observations suggest that depletion of Mop alters the organization of intracellular membrane compartments that support DCV replication. Whether the altered distribution of VP1 reflects changes in viral replication organelles, endosomal compartments or autophagic membranes remains to be determined.

One notable finding of this study is the striking contrast between cultured cells and adult flies. Whereas depletion of *mop* increased DCV replication in S2 cells, knockdown of *mop* in the adult fat body reduced systemic infection and significantly improved survival. At first glance these observations appear contradictory. However, the two experimental systems differ fundamentally. S2 cells measure the cell-autonomous function of Mop within infected cells, whereas fat body-specific knockdown influences the physiology of an organism in which the fat body functions as the principal organ for humoral immunity and metabolism. The fat body continuously secretes immune effectors, lipid carriers and signaling molecules into the hemolymph, thereby regulating systemic immunity and metabolism (Zhang, Tettamanti et al. 2021, Krejčová and Bajgar 2025). We therefore propose that the reduced susceptibility of mop KD flies may reflect changes in the systemic environment rather than the direct antiviral activity of Mop within individual infected cells.

Previous study demonstrated that Mop is required for efficient Toll signaling by promoting activation of antimicrobial peptide gene expression (Huang, Chen et al. 2010). This positive role of Mop for Toll signaling is preserved between S2 cells and adult fruit fly. It is also consistent with our observation that depletion of *mop* enhanced DCV infection in S2 cells. Although Toll signaling is classically associated with antibacterial and antifungal immunity, increasing evidence indicates that it also contributes to antiviral defense in *Drosophila*, either directly or through crosstalk with other immune pathways (Zambon, Nandakumar et al. 2005, Ferreira, Naylor et al. 2014). Loss of Mop may therefore weaken cell-autonomous antiviral defenses by compromising Toll-dependent immune responses, thereby allowing increased DCV replication in cultured S2 cells. In contrast, our *in vivo* analyses indicate that the reduced DCV infection observed in mop KD flies cannot be explained by enhanced activation of either the STING or JAK–STAT pathways, as neither *Srg3* nor *TotM* expression was elevated following *mop* knockdown. These observations suggest that the systemic antiviral phenotype is unlikely to arise from constitutive activation of these canonical antiviral signaling pathways and instead reflects a distinct physiological consequence of fat body-specific Mop depletion.

Transcriptomic analysis further supports this interpretation. Among the genes affected by *mop* knockdown, those involved in cell-cell junctions were preferentially upregulated, whereas genes associated with the Turandot family and ribosome biogenesis were downregulated. The decreased expression of *TotA*, *TotB*, *TotC* and *TotM* is unlikely to represent suppression of immunity because Turandot genes are induced by diverse forms of physiological stress and tissue damage in addition to infection (Rommelaere, Carboni et al. 2024). Their reduced expression may therefore simply reflect diminished tissue damage resulting from lower viral replication. Likewise, reduced expression of ribosome biogenesis genes may contribute to limiting viral protein synthesis (Cherry, Doukas et al. 2005), although whether this is a cause or consequence of reduced infection remains unclear. Proteomic analysis identified Ref(2)P and AGO2 as two of the most interesting proteins increased in mop KD flies. Ref(2)P is the *Drosophila* orthologue of mammalian p62/SQSTM1 and functions both as a selective autophagy receptor and as a restriction factor against sigma virus (Contamine, Petitjean et al. 1989, Avila, Silverman et al. 2002, Bartlett, Isakson et al. 2011). AGO2 is the central effector of antiviral RNA interference, the major antiviral defense pathway in insect cells (van Rij, Saleh et al. 2006). Increased abundance of these proteins is consistent with disruption of endolysosomal trafficking following Mop depletion. Because Mop is required for efficient lysosomal homeostasis, impaired trafficking may lead to altered turnover of Ref(2)P and AGO2. Alternatively, increased abundance of these proteins may represent an adaptive response that enhances antiviral defense. At present, our data do not distinguish between these possibilities. Determining whether autophagic flux and RNAi activity are functionally enhanced in mop KD flies will be an important objective for future studies.

Our study raises the broader possibility that Mop regulates the composition of factors secreted from the fat body into the hemolymph. Because the fat body is the major source of circulating immune proteins, lipoproteins and metabolic regulators, perturbation of endosomal trafficking could alter systemic signaling without necessarily changing canonical antiviral pathways such as STING or JAK–STAT. Such changes may influence the susceptibility of multiple tissues to DCV infection following systemic inoculation. This hypothesis also provides a plausible explanation for the opposite phenotypes observed in cultured cells and adult flies, in which the systemic consequences of altered fat body physiology override the cell-autonomous antiviral function of Mop. Identification of the circulating factors altered by Mop depletion, for example through quantitative hemolymph proteomics or transfer experiments using hemolymph from mop KD flies, will provide an important direction for future investigation.

In conclusion, our study identifies Mop as a previously unrecognized regulator of DCV infection in *Drosophila*. While Mop functions as a cell-autonomous restriction factor in cultured S2 cells, depletion of Mop in the adult fat body unexpectedly confers systemic resistance to DCV infection. These findings highlight the fundamentally different contributions of intracellular antiviral mechanisms and organismal physiological regulation during viral infection. More broadly, our results suggest that endosomal trafficking proteins can influence antiviral defense not only by acting within infected cells but also by modulating systemic homeostasis that determines host susceptibility to infection.

## CRediT authorship contribution statement

Tatsuhiko Kadowaki conceived and designed research strategy and wrote the paper. Jiaxin Liu, Kexin Li, Qinyi Liang, Yunfun Huang, and Xuye Yuan performed the experiments.

## Acknowledgements

This work was supported by Jinji Lake Double Hundred Talents Programme to TK. The funder had no role in study design, data collection and analysis, decision to publish, or preparation of the manuscript.

## Ethics statement

Not applicable

## Availability of data and materials

The RNA-seq data have been deposited in the NCBI Sequence Read Archive under accession number PRJNA1498194. The mass spectrometry proteomics data have been deposited to the ProteomeXchange Consortium via the PRIDE partner repository with the dataset identifier PXD081508. All other data is included in the manuscript and Supplementary material.

## Appendix A. Supplementary material

Supplementary Dataset 1

List of primers used in this study.

Supplementary Dataset 2

List of differentially expressed mRNAs in fat bodies isolated from DCV-infected *mop*-knockdown flies.

Supplementary Dataset 3

Gene Ontology (GO) terms enriched among the differentially expressed mRNAs listed in Supplementary Dataset 2.

Supplementary Dataset 4

List of differentially expressed proteins in fat bodies isolated from DCV-infected *mop*-knockdown flies.

